# Absenteeism and indirect costs during the year following the diagnosis of an operable breast cancer

**DOI:** 10.1101/514190

**Authors:** Clement Ferrier, Clémence Thebaut, Pierre Levy, Sandrine Baffert, Bernard Asselain, Roman Rouzier, Delphine Hequet

## Abstract

**Introduction:** The consequences of disease on work for individual patients as well as the consequences of absenteeism from work are subjects of interest for decision-makers.

**Methods:** We analyzed duration of absenteeism and related indirect costs for patients with a paid job during the year following diagnosis of early-stage breast cancer in the prospective OPTISOINS01 cohort. A human capital and friction costs approaches were considered for evaluation of lost working days. For this analysis, the friction period was estimated from recent French data. Statistical analysis included simple and multiple linear regression to identify determinants of absenteeism and indirect costs.

**Results:** 93% of patients had at least one period of sick leave, with an average of 2 periods of sick leave and a mean total duration of 186 days. 24% of patients returned to work part-time after an average sick leave of 114 days (i.e. 41 LWD). Estimated indirect costs were €22,722.00 and €7,724.00 per patient, for the human capital and friction cost approaches, respectively. In the multiple linear regression model, factors associated with absenteeism were: invasive tumor (p=.043), mastectomy (p=.038), redo surgery (p=.002), chemotherapy (p=.027), being a manager (p=.025) or a craftsman (p=.005).

**Conclusion:** Breast cancer is associated with long periods of absenteeism during the year following diagnosis, but almost all patients were able to return to work. Major differences in the results were observed between the friction cost and human capital approaches, highlighting the importance of considering both approaches in such studies.

## Introduction

Breast cancer (BC) is the most common cancer in women in France with an estimated incidence of 54,062 new cases in 2015 (standardized incidence rate of 97.4 cases per 100,000 women per year). Costs associated with health care interventions are usually divided between direct costs, referring to resource utilization, and indirect costs, representing loss of productivity or the time cost due to illness. Various Health Technology Assessment (HTA) agencies have developed guidelines identifying the cost components to be included in economic evaluation, and generally recommending exclusion of indirect costs for various reasons. Alternatively, the French Health Authority (HAS) has proposed considering the production costs of health care interventions, including the standard direct costs and the time cost of delivery of care to patients, but excluding the time cost due to the underlying condition [1].

BC generates major direct costs for health care systems. A study conducted in France on the national health insurance database concluded that BC was the most expensive type of cancer, representing 18.5% of all cancer expenditure [2]. In the US, the estimated lifetime treatment costs ranged from $20,000 to $100,000 [3]. Moreover, since BC is mostly prevalent in women of working age, it induces considerable indirect costs, which should be taken into account to ensure a more comprehensive approach to this disease from a societal perspective.

The methodology of indirect cost measurement raises a number of issues, especially estimation of lost productivity related to the time lost due to illness. The human capital (HC) approach, defined by Rice, evaluates the entire time lost, regardless of the duration of time off work [4]. In contrast, the friction costs (FC) approach assumes that indirect costs only occur during a limited period before the employer company adjusts to the worker’s absence, defined as a friction period [5]. Although both of these approaches can provide useful information, the two approaches provide estimates that can differ dramatically, especially when considering a lifetime horizon [6-8].

Data concerning absenteeism and indirect costs are useful for decision-makers when considering cost-of-illness studies or model-based economic evaluations on innovative treatments or screening strategies. Few published studies have focused on the indirect costs of BC and absenteeism, with estimates ranging from $8,068 [9] to $21,086 per patient [10], highlighting the importance of the methodology used and the country in which the study is conducted. In France, only one study has reported the indirect costs associated with adjuvant chemotherapy in BC [11]. Only limited data are also available concerning the individual characteristics driving the level of indirect costs. It may be justified to maintain employment or promote early return to work in order to decrease the economic burden of the disease from a societal perspective, but also to limit the impact of the disease on the patient’s personal life (social environment, long-term career goals, etc.). In some countries, such as France, return to work is considered to be a public health priority [12-13] and could be promoted by means of multidisciplinary interventions and less toxic innovative treatments [14].

The objective of this study was to describe the indirect costs of absenteeism and their determinants in the first year following the diagnosis of early BC in a French population-based prospective cohort study, using both the HC and FC approaches.

## Material and methods

### Population

We analyzed indirect costs in early BC during the year following diagnosis. This study was part of a global research on BC pathways and burden of disease of the French multicenter OPTISOINS01 study. The design of this prospective trial has been previously described [15]. Female patients with histologically confirmed, previously untreated and primarily operable BC (exclusion of metastatic, locally advanced or inflammatory BC as defined by the AJCC) were included in this cohort by 8 centers (three University hospitals, four local hospitals, one comprehensive cancer center) in three departments of the Ile-de-France region between 2014 and 2016. Patients were prospectively followed for one year with data collection concerning three work packages: 1/resource utilization and costs of pathways, 2/patient satisfaction and work reintegration, 3/quality, coordination and access to innovation. Individual social and economic characteristics were also recorded at the beginning of the study. Informed consent was obtained from all individual participants included in the study.

All patients of the initial OPTISOINS01 cohort reporting a paid job at the time of the diagnosis were included in the present study. Patients with missing data on wage or absenteeism were excluded.

All components of absenteeism were included as part of indirect costs: days of sick leave, part-time return to work, early retirement, mortality. Indirect costs related to presenteeism and unpaid work were excluded from the scope of this study. A specific questionnaire in the second work package was used to assess time lost due to the disease during the year of the survey: dates of work and absence from work during treatment, work arrangements, on-shift status (e.g., recognition of disability at work, applications for disability allowance, retirement, and layoff). Indirect costs for relatives were not taken into account.

### Methods used to calculate indirect costs

Periods of absenteeism were considered by both the HC and FC approaches, applied separately. In the HC approach, we assumed that indirect costs were generated during the entire period of absenteeism [4]. According to the FC approach [6], after a friction period, the level of productivity was restored due to replacement of the sick worker, incurring no indirect costs. However, medium-term global macroeconomic consequences may arise in an international competitive labor market with an impact on macroeconomic indicators. During the friction period and as a result of various parameters (diminishing returns to labor, internal labor reserve within firms, and delaying work after the period of absence), production losses are lower than estimates based on the HC approach. With the FC approach, indirect costs were restricted to the friction period and a friction coefficient was used to reduce the value of lost production. Medium-term macroeconomic consequences were assumed to be insignificant for this cohort.

According to Koopmanschap’s method, job vacancy duration estimates (i.e. length of recruitment processes) increased by a 30-day time lag (for the decision to recruit and the time between recruitment and the first working day) were used as a proxy for the duration of the friction period [6]. As job vacancy duration depends on job categories, job vacancy duration estimates were stratified for this factor (managers vs. other job categories). Data concerning job vacancy durations have been rarely reported in the French literature (including the grey literature). The *OFER* survey, conducted in 2005 by the labor department, reported mean job vacancy durations of 56 days for managers and 28 days for other job categories [16]. In 2017, *Pole-emploi* job offers were filled after an average of 56 days for managers and 36 days for other categories [17]. Consequently, we assumed a friction period of 86 days for managers and 62 days for other job categories. A value of 0.8, initially computed by Koopmanschap et al. (using the estimated elasticity for annual labor time versus labor productivity), which has never been subsequently updated, was used for the friction coefficient [6].

Periods of absenteeism, including days off (weekends), were recorded in the patient’s questionnaire and converted into Lost Working Days (LWD) for the analysis: five LWD for 7 sick days, 2.5 LWD for 7 days of part-time return to work, 251 LWD for death or early retirement (considering 5 working days per week during a year). LWD, representing lost productivity, were estimated in euros based on individual daily wages plus employers’ and employees’ social welfare contributions (computed by an online tool [18]). Wages, expressed as monthly net income, were self-reported by the patients. When multiple periods of absenteeism, interspersed by periods of work, were recorded in the FC approach, an entire new friction period was assumed for each period of absenteeism.

### Statistical analysis

Various simple and multiple linear regression models were performed using Stata\IC 14.0 software (StataCorp, College Station, Texas). In the first model, the independent variable was the number of LWD estimated by the HC approach. In the second model, the independent variable was the amount of indirect costs. As HC and FC approaches result in different values, two distinct statistical analyses were performed. The following dependent variables were integrated in a simple linear regression: marital status, job category, cancer histology, surgical treatment, adjuvant therapy for BC. Factors correlated with the independent variable (with a p-value<.1) were integrated in the multiple linear regression. A p-value < .05 was considered significant.

A univariate sensitivity analysis was conducted on the parameters estimated by the FC approach (duration of the friction period, friction coefficient) by applying ±20% variation and a Tornado diagram was constructed to illustrate the effect on indirect costs. This analysis was also conducted for subgroups: managers and other job categories.

To facilitate the use of our data in other health economics analyses, average indirect costs according to subgroups defined by care pathway and job category were also estimated.

This study was approved by the French National ethics committee (CCTIRS Authorization n°14.602 and CNIL DR-2014-167) covering the research conducted at all participating hospitals. This study was registered with ClinicalTrials.gov (Identifier: NCT02813317).

## Results

Six hundred four of the 617 screened patients were included in the OPTISOINS01 cohort. Some patients were excluded from the study either due to the absence of a paid job during the study period (n=307) or due to missing data (wages for 42 patients and length of absenteeism for 87 patients). A total of 168 patients were included in the analysis (Figure 1). Patient characteristics are presented in Table 1.

**Figure 1.**
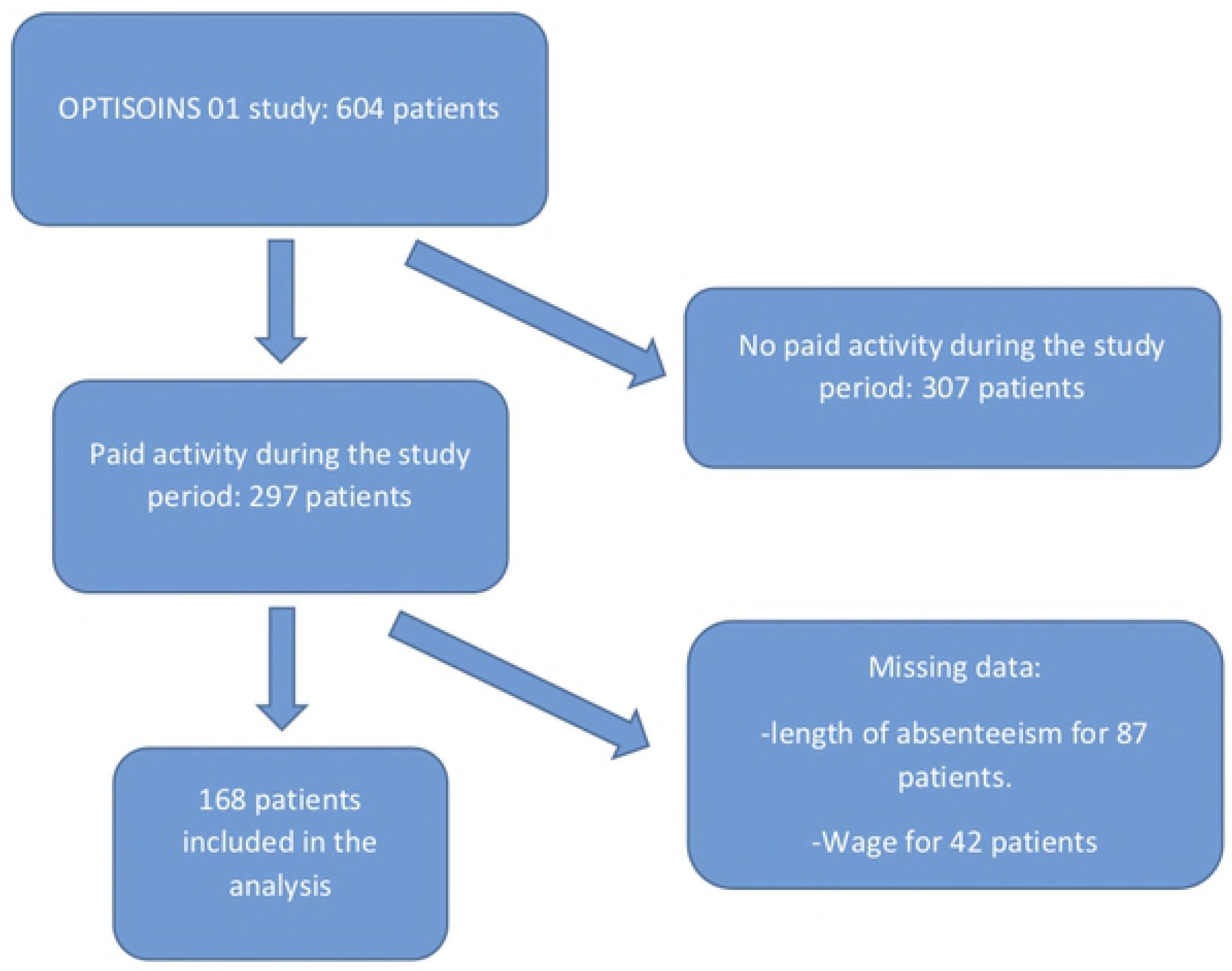
Flow-chart.

**Table 1.** Characteristics of the sample.

|  |  |
| --- | --- |
| **Age<sup>1</sup> (years)** | 50.1 (±7.4) |
| **Marital status** |  |
| -Single | 31 (18.5%) |
| -Married | 106 (63.1%) |
| -Divorced | 28 (16.7%) |
| -Widow | 1 (0.6%) |
| -Missing data | 2 (1.2%) |
| **Educational attainment** |  |
| -Elementary | 10 (6%) |
| -Middle school | 36 (21.4%) |
| -High school | 24 (14.3%) |
| -Post High school | 98 (58.3%) |
| **Job category** |  |
| -Employee | 70 (41.7%) |
| -Manager | 70 (41.7%) |
| -Temporary worker | 20 (11.9%) |
| -Craftsman | 5 (3%) |
| -Missing data | 3 (1.8%) |
| **Monthly wage<sup>1</sup> (€)** | 2,214 (±761) |
| **Mode of diagnosis** |  |
| -Screening | 69 (41.1%) |
| -Other | 99 (58.9%) |
| **Histology** |  |
| -In situ | 21 (12.5%) |
| -Invasive | 146 (86.9%) |
| -Missing data | 1 (0.6%) |
| **Surgery** |  |
| -Lumpectomy | 129 (76.8%) |
| -Mastectomy | 39 (23.2%) |
| -Sentinel lymph node | 121 (72%) |
| -Axillary lymph node dissection | 33 (19.7%) |
| -No lymph node surgery | 14 (8.3%) |
| **Number of surgical procedures** |  |
| -1 | 129 (76.8%) |
| -2 | 36 (21.4%) |
| -3 | 3 (1.8%) |
| **Adjuvant radiation therapy** | 153 (91.1%) |
| **Adjuvant chemotherapy** | 80 (47.6%) |
| **Adjuvant endocrine therapy** | 121 (72%) |
| **Sick leave** |  |
|  | 156 (92.9%) |
| -Proportion with a least one | 2 |
| period | 173 ($\pm 155$) |
| -Number of periods<sup>1</sup> | 186 ($\pm 153$) |
| -Number of days<sup>1</sup> |  |
| -Number of days if at least one |  |
| period<sup>1</sup> |  |
| **Part-time return to work** |  |
| -Proportion | 40 (23.8%) |
| -Number of periods<sup>1</sup> | 1 |
| -Number of days<sup>1</sup> | 27 ($\pm 63$) |
| -Number of days if part-time return | 114 ($\pm 83$) |
| to work<sup>1</sup> |  |
| *1: mean (+/-standard deviation).* |  |

Eleven patients reported no periods of absenteeism and 156 patients (93%) reported at least one period of sick leave, with an average of 2 periods and 186 days of sick leave (i.e. 133 LWD). Forty patients (24%) reported part-time return to work after an average of 114 days of sick leave (i.e. 41 LWD). Five patients requested early retirement, but procedures were still pending at the end of follow-up. No death was declared during the study period. On average, managers declared fewer sick leave days than other job categories (146 and 172, respectively), but longer periods of part-time return to work (33 and 24 days, respectively). A total of 20,973 LWD (mean: 125 LWD per patient) was estimated by the HC approach and a total of 8,297 LWD (mean: 49 LWD per patient) was estimated by the FC approach. No significant difference in the number of LWD estimated by the two approaches was observed for 66 patients (38%). Patients in this subgroup presented early-stage disease with significantly lower rates of mastectomy (11%), axillary lymph node dissection (5%) and chemotherapy (27%) than the overall sample.

Consequently, the total indirect costs in the cohort estimated by the HC approach were threefold higher than those estimated by the FC approach: €3,817,000.00 vs. €1,298,000.00, representing €22,722.00 and €7,724.00 of indirect costs per patient during the first year after diagnosis, respectively. In both approaches, the main driver of lost productivity and indirect costs was the number of days of sick leave, representing 90% of the total cost. Part-time return to work accounted for only10% of total costs (Figure 2). Indirect costs were higher for managers than for other job categories: €11,044/patient vs. €5,353/patient with the FC approach, despite fewer sick leave days for managers. This result can be explained by the higher incomes of managers.

**Figure 2.**
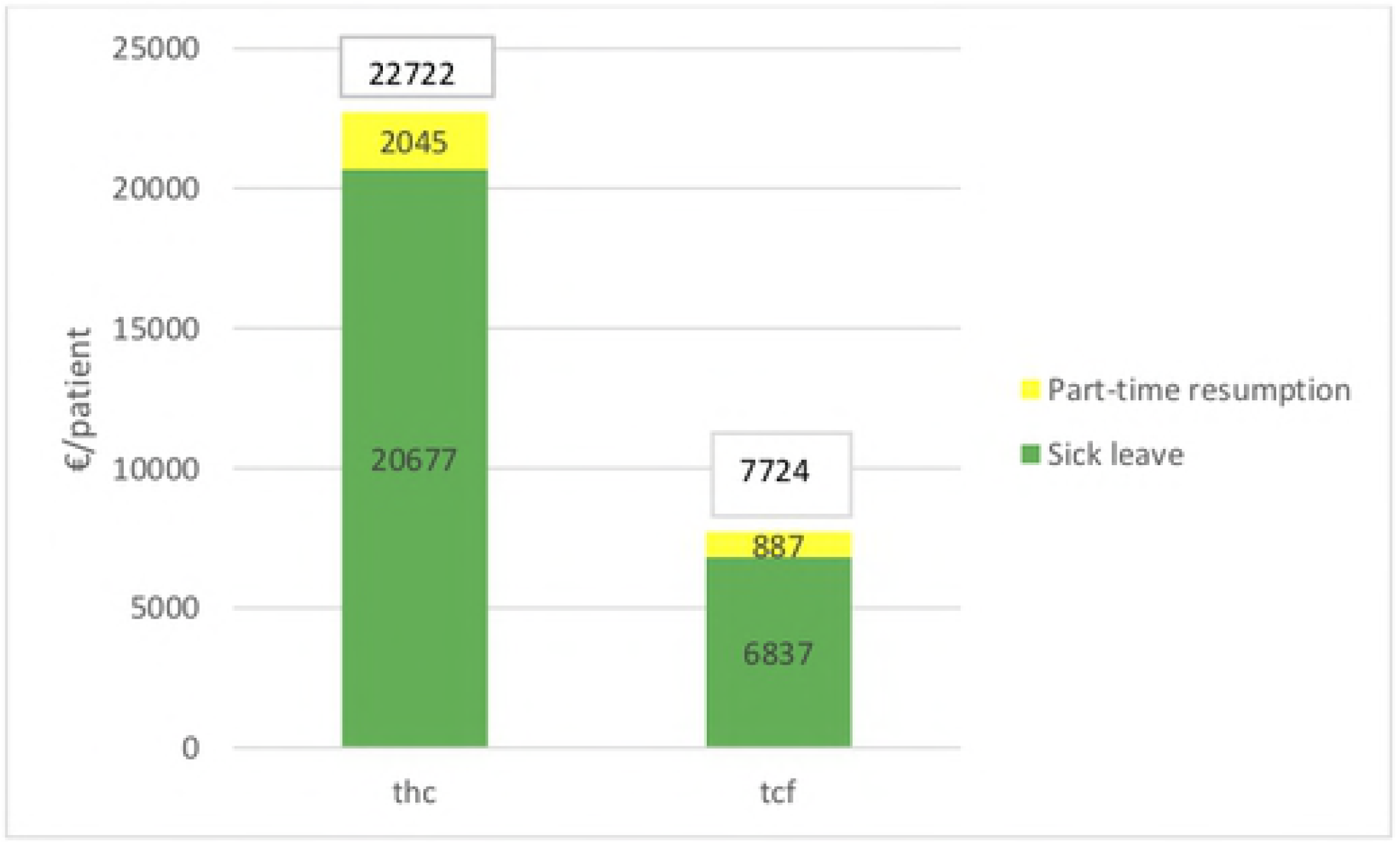
Indirect costs per patient.

The following factors were associated with the number of LWD in the multiple linear regression model: invasive cancer (+65 LWD, p=.043), mastectomy (+40 LWD, p=.038), redo surgery (+48LWD, p=.002), chemotherapy (+35 LWD, p=.027), being a manager (-33LWD, p=.025) or a craftsman (-113LWD, p=.005) (compared to being a salaried employee). According to both approaches, in the simple linear regression model, individual indirect costs were correlated with invasive cancer, treatment characteristics (mastectomy, axillary lymph node dissection, chemotherapy) and socioeconomic characteristics (highly educated vs. poorly educated, being manager vs. being a salaried employee) (see Table 2. For details and differences between HC and FC approach). According to the HC approach, the following factors were significantly associated with indirect costs in the multiple linear regression model: mastectomy (p=.046), redo surgery (p=.003) and being a craftsman (p=.023). Indirect costs in the model were increased by being a manager, having an invasive tumor or having received chemotherapy, but with a lower degree of statistical significance. According to the FC approach, being a manager was the only significant factor associated with indirect costs in the multiple linear regression model (p<.001) (Table 2).

**Table 2.** Regression models on indirect costs.

|  | HC approach-SLR |  | HC approach-MLR |  | FC approach-SLR |  | FC approach-MLR |  |
| --- | --- | --- | --- | --- | --- | --- | --- | --- |
|  | Coef.<br>[95%CI] | p-value | Coef.<br>[95%CI] | p-value | Coef.<br>[95%CI] | p-value | Coef.<br>[95%CI] | p-value |
| Age | -18.7 | .928 | - | - | 21.4 | .32 | - | - |
| Marital status |  |  | - | - |  |  | - | - |
| -Single |  |  |  |  |  |  |  |  |
| -Married | 1515 | .71 |  |  | 1152 | .384 |  |  |
| -Divorced | 3770 | .471 |  |  | 1076 | .524 |  |  |
| Educational attainment |  |  | - | - |  |  | - | - |
| -post high school | -4867 | .462 |  |  | -5458 | **.08** |  |  |
| -elementary | -6785 | **.082** |  |  | -4160 | **.001** |  |  |
| -middle school | 1462 | .747 |  |  | -1029 | .464 |  |  |
| -high school |  |  |  |  |  |  |  |  |
| Job category |  |  |  |  |  |  |  |  |
| -Salaried employee |  |  |  |  |  |  |  |  |
| -Manager | 5573 | **.096** | 4838 | .117 | 5694 | **<.001** | 5688 | **<.001** |
| -Temporary | -1726 | .73 | -618 | .894 | 1505 | .305 | 1838 | .212 |
| -Craftsman | -15819 | **.085** | -19231 | **.023** | -3760 | .161 | -4248 | .111 |
| Diagnosed by<br>screening | -898.7 | .775 | - | - | -858 | .394 | - | - |
| Invasive<br>nature | 13849 | **.003** | 11676 | **.086** | 2743 | **.067** | 1573 | .462 |
| Breast surgery<br>-lumpectomy<br>-mastectomy | 13549 | **<.001** | 8183 | **.046** | 2489 | **.033** | 859 | .505 |
| Axillary<br>surgery<br>-sentinel<br>lymph node<br>-no dissection<br>-complete<br>dissection | -6692<br>14023 | .216<br>**<.001** | 3152<br>5271 | .676<br>.215 | -1591<br>3122 | .371<br>**.012** | 458<br>2297 | .847<br>**.088** |
| Number of<br>surgical<br>procedures | 7710 | **.018** | 9624 | **.003** | 1465 | .162 | 1652 | .102 |
| Chemotherapy | 11300 | **<.001** | 5245 | **.113** | 1760 | **.075** | 420 | .687 |
| Radiotherapy | -9240 | **.087** | -4003 | .465 | -1756 | .312 | -1282 | .459 |
| Endocrine<br>therapy | 5743 | **.094** | 1837 | .624 | 1135 | .303 | 732 | .537 |

*SLR: Simple Linear Regression. MLR: Multiple Linear Regression. HC: Human Capital. FC: Friction costs.*

In the univariate sensitivity analysis based on the FC approach, the variation of friction coefficient (benchmark: 0.8; +/-20%) had the greatest impact on indirect costs, which were increased from €6,179/patient to €9,269/patient (basis: €7,724/patient) (Figure 3). In the subgroup analysis, indirect costs were much more sensitive to variations of estimated parameters for managers than for other job categories.

**Figure 3.**
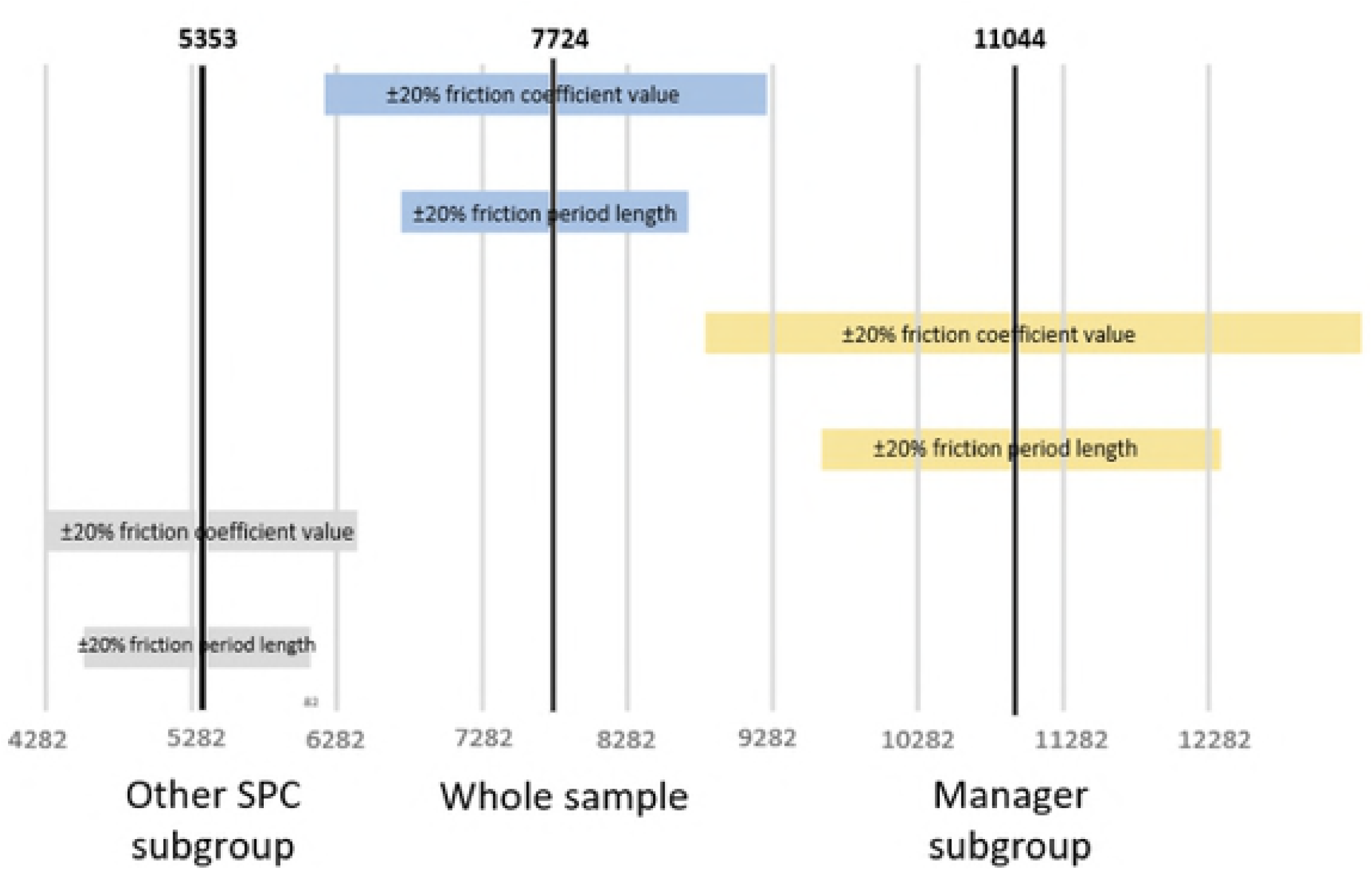
Tornado diagram for univariate sensitivity analysis.

Tables with estimated average indirect costs by care pathway and job category are presented in supplementary data.

## Discussion

Direct costs associated with breast cancer have been extensively documented in the literature, but the determinants of indirect costs are less well known. Studying the OPTISOINS cohort, Arfi et al. [19] estimated the average daily sick leave allowances (i.e. indirect medical costs) to be about €8,841 per patient for the first year after diagnosis. In the same cohort, we showed that BC is often associated with periods of sick leave and part-time return to work during the year following diagnosis, resulting in considerable indirect costs even in the absence of mortality. However, the use of two different approaches resulted in discordant estimated indirect costs per patient: €22,722.00 for the HC approach *vs.* €7,724.00 for the FC approach. The following factors were correlated with the number of LWD and the sum of indirect costs in the HC approach: invasive tumor, mastectomy, redo surgery, chemotherapy, being a manager or a craftsman. Indirect costs estimated by the FC approach were mostly driven by job category, as managers had higher wages and longer friction periods.

The mean age at the diagnosis of BC corresponds to the mean age of retirement in industrialized countries (62 years in France) [20]. Moreover, retirement age is currently increasing in many countries, as a result of increasing life expectancy [21]. Absenteeism, indirect costs and, more generally, the work consequences associated with BC constitute major concerns for decision-makers. In the initial OPTISOINS01 cohort, one-half of patients had a paid job during the study period. These patients spent an average of almost six months on sick leave during the year after diagnosis and 24% of patients returned to work part-time for a mean period of 114 days. A mean of 125 LWD per patient was recorded and 26 patients were absent from work for the entire year of analysis, highlighting the impact of BC on paid employment. Only a few studies have previously reported these data. Two US retrospective cohort studies found that working patients with BC had significantly longer periods of absenteeism compared to “control” patients (without BC) [9,22]. However, they observed a shorter period of absenteeism in the year following diagnosis when using an HC approach: about 10 days of sick leave and between 25 and 45 days of short-term disability (according to the severity of the disease). In Sweden, patients reported on average 271 working hours lost during the 3 months preceding the interview during the first year after a diagnosis of primary BC [10]. In line with our results, mortality and early retirement were rare during the first year following diagnosis.

The present study is the first prospective cohort study focusing on the cost of BC-related absenteeism. The large initial sample size of OPTISOINS01 allowed us to exclude working patients with missing data, which could have affected the quality of the analysis. Due to the geographical design of the study, most patients in this cohort lived near Paris and may have presented different socio-economic characteristics to those of the national population. For example, the proportion of managers in this sample was larger than in the French working population (41.6% vs. 17.1%), and the average wage was also higher (€2,214 vs. €1,926) [23]. The timeframe was limited to one year after diagnosis, but most indirect costs related to BC occur during the first year, except when the disease progresses to metastatic stage or recurrence [10,24].

This study only considered indirect costs due to absenteeism, but there is a growing interest in presenteeism, i.e. workers who have returned to work, but with reduced capacity and productivity due to illness, and unpaid work, which generate indirect costs [25]. Appropriate tools (iMTA Productivity Cost Questionnaire [26, Valuation of Lost Productivity [27]) must be used to investigate these phenomena, but were not used in the present cohort. This component was therefore excluded from this analysis. Although a few studies have suggested substantial indirect costs associated with presenteeism and unpaid work [24,28,29], no guidelines concerning the integration of these factors have been established.

Published studies are generally based on a single method for calculation of lost productivity, usually the HC approach. When using the FC approach, authors use friction periods initially estimated by Koopmanschap et al. (2.8-3.2 months [6]), or other values but with no justification of their sources [30]. However, Koopmanschap’s estimation is based on outdated data (1988-1990) restricted to the Dutch labor market. The time to fill job vacancies (i.e. the length of recruitment processes) increased by a time lag of four weeks (decision to recruit and time between recruitment and the first working day) was used to estimate the friction period. The duration of the friction period is therefore strongly correlated with the unemployment rate and the economic environment and varies over time and across countries. The duration of the friction period must also be stratified according to job category, as the recruitment process is longer for managers than for employees [31]. In the absence of guidelines, we preferred to use both approaches and used French grey literature sources to estimate the duration of friction periods according to Koopmanschap’s method. Our estimation of the duration of friction periods was similar to that proposed by Koopmanschap for managers, but much shorter for other job categories (62 vs. 90 days), highlighting the importance of using appropriate data for the FC approach. Nevertheless, our sensitivity analysis focused on the impact of the estimated friction coefficient on the sum of indirect costs. In order to improve the robustness of our results, we would need to perform an estimation for France, which would require microeconomic data that unfortunately were not available for the companies concerned. Friction period duration and friction coefficient values should therefore be regularly estimated and updated to facilitate application of the FC approach. Taieb et al propose a new approach to compare the HC and FC methods which are used to measure health-related production losses to produce guidelines on which method should be used rather than trying to establish the superiority of one method over the other [32]. They showed that the choice between the HC and the FC should be based on two main criteria: the type of absence from work under consideration (sickness absence or death) and the type of economic evaluation used (Cost-Effectiveness, Cost-Utility etc.). These guidelines are elaborated so as to avoid underestimation and double counting of the costs of illness.

As indicated above, the need to account for indirect costs related to productivity losses in economic studies has not been clearly established. Health Technology Assessment agencies provide different guidelines. Guidelines focusing on the payer perspective recommend the exclusion of these costs from the evaluation [33,34]. In contrast, other agencies adopting a collective or societal perspective, including the World Health Organization, propose the inclusion of indirect costs, sometimes in a separate analysis [35,37]. Apart from the methodological challenge of measuring indirect costs, this approach also raises a number of ethical issues [38], as, if decision-makers use these studies to plan resource allocations, working age people would be favored over young or elderly, and other unemployed populations. Several authors have also discussed the risk of double counting in cost-effectiveness evaluation when productivity losses are included. Do individuals take the potential impact on their income and career into account when assessing the impact of their disease on quality of life, included in the denominator of the incremental cost-effectiveness ratio (ICER) [39,40]. Separate analysis of direct and indirect costs can help to guide decision-makers in relation to these scientific and equity issues in healthcare resource utilization.

The relationship between disease and work also needs to be studied in more detail in order to limit the consequences of illness on work. The low rate of early retirement and the absence of dismissal in our cohort should reassure BC patients concerning their future capacity to work and should encourage them to return to work part-time when necessary. Another study showed good readjustment to the workplace for patients with BC [41]. In agreement with many authors, we therefore believe it is crucial to report absenteeism, productivity losses and indirect costs. Moreover, accurate data on the type and duration of absenteeism are essential, particularly for economic studies based on simulated models and using a societal perspective.

## Acknowledgment

This study was supported by a grant from the French National Cancer Institute, dedicated to economic studies of innovative techniques (PRME-K-2013).

## Figure legends

Figure 2: Tch: Human Capital Approach. Tcf: Friction Costs Approach.

Figure 3: X-axis: cost (€) / patient. SPC: job category. For the subgroup analysis, the variation is applied only for the patients in the subgroup.

